# Differences in apple fruit shape are independent of fruit size

**DOI:** 10.1101/2025.01.01.631017

**Authors:** Kylie DeViller, Daniel H. Chitwood, Sean Myles, Mao Li, Zoë Migicovsky

## Abstract

**Premise:** Fruit quality is crucial in breeding new apple accessions. Before tasting, consumers assess freshness and flavour based on the physical appearance of fruit. Understanding how fruit quality traits such as shape and size vary across diverse apples provides a foundation for future breeding efforts.

**Methods:** We analyzed images of 5724 apples representing 743 different trees and 534 unique accessions from Canada’s Apple Biodiversity Collection to quantify variation in fruit shape and size. To achieve this, we used a pseudo-landmarking approach paired with traditional linear measurements including length, width, area, solidity, and aspect ratio. We also incorporated previously collected fruit weight measurements from the same trees.

**Results:** Using a comprehensive measure of shape, we determined that the primary source of variation in apple fruit shape was the width to length (aspect) ratio of the fruit. This variation was not significantly correlated with differences in fruit size from its area and harvest weight.

**Conclusions:** Our findings indicate that two critical aspects of morphological variation in apple—fruit shape and size—are independent, suggesting it is possible to select for a diverse range of fruit shapes while maintaining a consistent and marketable size.

## Introduction

External appearance is the first metric consumers use to assess the quality of fresh fruit, which makes shape, size, and colour critical breeding targets (Kays, 1999). Appearance also serves as a visual cue that can trigger appetite and memories of previous eating experiences (Berthoud & Morrison, 2008). Deviations in fruit appearance from consumer norms can result in the perception that fruit is inferior in terms of flavour or juiciness (Delwiche, 2012). While some abnormalities in shape are acceptable to consumers, fruit that are extremely oddly shaped are not, and these are frequently discarded during harvesting or packaging (Kays, 1999; Loebnitz et al., 2015).

Fruit shape is a critical consideration for apple breeders, as they must consider uniqueness in terms of flavour, texture, and appearance (Brown & Maloney, 2005). Consumers use fruit shape to assist in identifying accessions (Musacchi & Serra, 2018) and it must be included when registering new accessions (International Union for the Protection of New Varieties of Plants, 2023). Traditionally, evaluations of fruit size were performed manually using callipers while fruit shape was subjectively scored based on human observation. Such manual and subjective approaches are time-consuming and laborious, and taking such measurements across a large number of fruit and trees remains prohibitive.

Recent methods have been developed for measuring fruit colour and shape using high-throughput and quantitative image analysis (Huang et al., 2020). With digital images, extraction of data can be automated, and images can be stored for future use. Two recent studies used image analysis for apple fruit shape. The first used Tomato Analyzer (Gonzalo et al., 2009) to extract measurements for 11 shape and 4 size attributes from fruit sections (Dujak et al., 2023) and the second used FruitPhenoBox (Kirchgessner et al., 2024), a custom built RGB camera system to automate data collection for fruit colour, shape, and size (Keller et al., 2024). Continuing improvement in quantitative image analysis of fruit shape can assist plant breeders in selecting for consumer-preferred traits while also improving knowledge of the boundaries of morphological variation (Chitwood & Mullins, 2022).

One approach for examining variation in morphology is the use of homologous landmarking based on shared morphology. This method has been widely applied in the study of leaf shape (Chitwood, Klein, et al., 2016; Chitwood, Rundell, et al., 2016; Chitwood & Otoni, 2017; Migicovsky et al., 2022). Homologous landmarking involves placing coordinate points at defined locations that are then used to capture overall shape (Ibragimov & Vrtovec, 2017; Webster & Sheets, 2010). These locations provide spatial information for mapping measurements and allow for comparison across samples based on shared homology (Webster & Sheets, 2010). However, this approach is not feasible when there are no, or few, shared landmarks. As an alternative, pseudo-landmarking can be used to estimate shape by placing equidistant points along the contour of an object (Bookstein, 1997; Ibragimov & Vrtovec, 2017). In contrast to human observation and subjective scoring of shape, both homologous landmarks and pseudo-landmarks can be paired with computational analyses to quantify overall morphological diversity in shape.

In this study we apply a pseudo-landmarking approach to 5724 fruit images taken from 534 unique apple accessions from Canada’s Apple Biodiversity Collection. We integrate these data with fruit weight measurements to improve our understanding of morphological diversity in apple. We determine that variation in the fruit width to length ratio is the primary driver of fruit shape diversity in apple but that this variation is size-independent. This work contributes to our understanding of apple morphology and provides a foundation for future efforts to understand the genetic basis underlying fruit shape to inform plant breeding.

## MATERIALS AND METHODS

### Data Collection

The fruit images used in this study were taken from Canada’s Apple Biodiversity Collection located at Agriculture and Agri-Food Canada’s (AAFC) Kentville Research and Development Centre (KRDC) in Nova Scotia, Canada. A comprehensive description of the design, maintenance, and harvest of the Canada’s Apple Biodiversity Collection can be found in Watts et al. (2021). Briefly, each of the 1119 unique accessions were grafted to a M.9 rootstock and planted in a randomized incomplete block design in 2013. In 2016, fruit were harvested, and images of each fruit (approximately 10 per tree) were taken using a custom-built light box. Although images were taken of both the top and side of the fruit, in this study only images of the side of the fruit were retained for a total of 5860 images. All images were uploaded to Dryad (URL to come). To focus on within-species variation, only images for trees identified as *Malus* X. *domestica* Borkh. according to Watts et al. (2021) were retained. A final dataset of 5724 images from 743 trees remained for downstream analyses.

### Image Processing and Morphometric Analysis

Images (Figure 1a) were colour corrected using the included colour card and then converted into binary scans using MATLAB® (The MathWorks Inc., 2018) following the methods described by Li et al. (2022). Binary scans were uploaded to accompany fruit images to Dryad (URL). After binary conversion, individual fruit were landmarked using ImageJ (Abràmoff et al., 2004). Two landmark points were placed in the middle of the top and bottom of the fruit (Figure 1b). If placement was unclear from the binary image, the original image was consulted to better identify the top and bottom. Coordinates for all images were merged into a table using R (v4.4.0) (R Core Team, 2024) in RStudio (RStudio Team, 2023). As all images were taken using the same device set up, an average pixels per cm measurement was taken across 10 images based on the included ruler. This value was used for scaling of size-related measurements described below.

**Figure 1.**
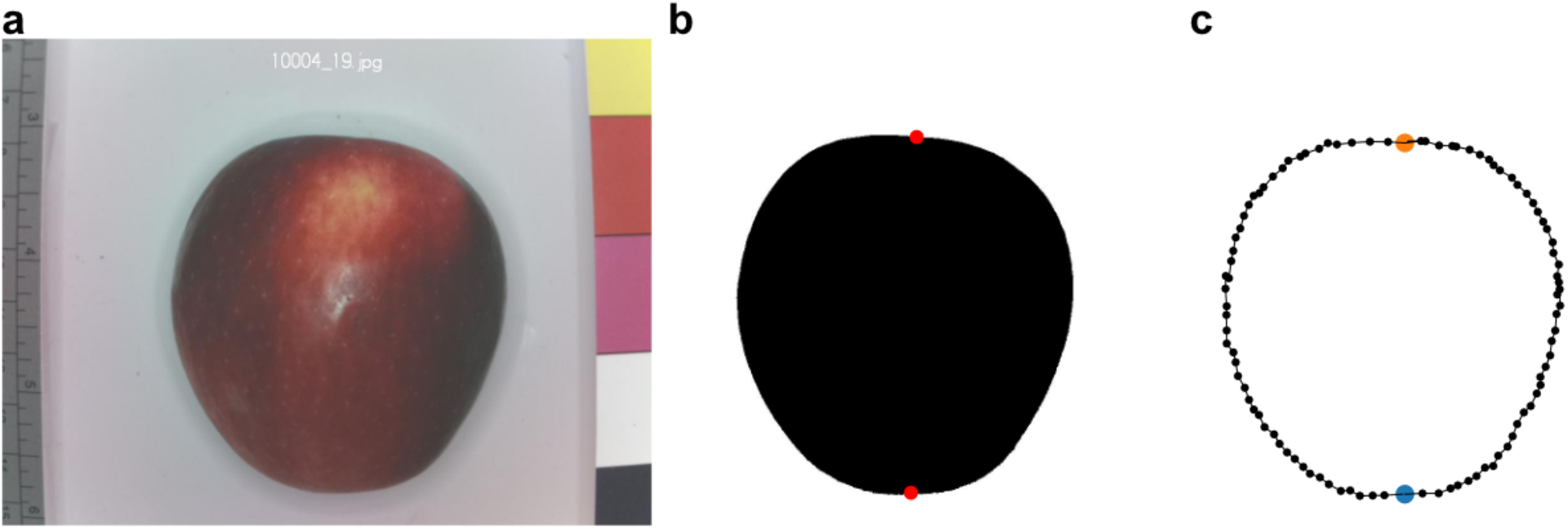
Methodology of landmarking. (a) Original colour corrected images that were used to extract the contour of the apple. (b) Following binary conversion, two landmarks were placed in the center of the top and the bottom of the fruit to provide reference points to align the fruit to each other in analysis. (c) 50 pseudo-landmarks were placed on each half of the fruit with one removed at the top for a total of 99 pseudo-landmarks.

Binary images were analyzed based on methods and custom functions in Python (v3.12) (Van Rossum & Drake, 2009) implemented in R (R Core Team, 2024; RStudio Team, 2023) using the reticulate R package (Ushey et al., 2024) with Quarto (Allaire et al., 2024). After reading in the binary images, the contour of the largest object (the fruit) was selected and interpolated using 1000 equidistantly placed pseudo-landmarks to facilitate identifying the landmark closest to the base. The top and bottom of the fruit, as indicated by the manually added landmarks, were then used so that the base index was zero. Next, each side of the fruit was interpolated using 50 pseudo-landmarks, equidistant on each side of the fruit. The total number of pseudo-landmarks was 2*50– 1, with the 1 being removed due to the overlap at the top, for a total of 99 pseudo-landmarks, with the contour starting and ending at the same point at the base. The segmented outline may appear that the landmarks are not perfectly equidistant due to the ridges along the outline. The fruit were then rotated based on the top and base landmarks to ensure that top and bottom coordinates align and that the top of the fruit was always upward facing. The pseudo-landmarks, representing the fruit, were scaled according to the pixels per cm value (Figure 1c).

Following the calculation of 99 pseudo-landmarks for each image, additional linear measurements were calculated based on the rotated and scaled images. Using the oriented fruit, width was calculated as the difference between the minimum and maximum of the x coordinate values, while length was calculated as the same difference for the y coordinates. Aspect ratio was calculated by dividing the width measurement by the length. The shoelace algorithm, which calculates the area of a polygon, was used to calculate the area of each apple fruit in cm^2^.

Solidity was calculated by dividing the area of each fruit by the convex hull area. The convex hull area was calculated separately and represents a contour that fully contains all landmarks. Thus, higher solidities indicate that there was less empty area between the convex hull and the contour of the fruit, while lower solidity values indicate a less circular fruit with more pronounced gaps between the convex hull and contour. Linear measurements were averaged across within-tree replicates then adjusted based on position in the orchard using a restricted maximum likelihood (REML) model, as described by Watts et al. (2021) using the R packages lme4 (Bates et al., 2024), pbkrtest (Halekoh & Højsgaard, 2024), lsmeans (Lenth, 2018), and lattice (Sarkar, 2008). This process reduced the dataset from 743, the number of individual trees sampled, to 534, the number of genetically unique accessions sampled.

### Statistical Analyses

Using 99 pseudo-landmarks for each image, a generalized Procrustes analysis (GPA) was performed using a custom function in Python. Following GPA, values were averaged across the replicates (approximately 10) within a tree. Next, principal components analysis (PCA) was performed. The first three principal components (PCs) were retained for analyses as they explained a total of 54.1% of the variation in shape and were the most visually distinguishable. PC values were REML-adjusted using the same method as described for linear measurements. PC values and linear measurements were merged with fruit weight measurements and release year data from Watts et al. (2021). Fruit weight was measured from the same fruit that were photographed. An accession’s “release year” refers to its year of commercialization, release to the public, or initial mention in historical records. All data included in this study such as linear measurements, GPA-adjusted landmarks, PCs, fruit weight, and year of release, are reported in Table S1.

To explore the relationships between fruit shape, size, and weight, we assessed correlation between all pairwise comparisons of the selected 10 traits using a Spearman’s rank correlation coefficient using the cor.test() function in the R stats package (R Core Team, 2024). We adjusted p-values using a Bonferroni correction based on the total number of comparisons (45). All pairwise comparisons were visualized using ggpairs (Schloerke et al., 2024; Wickham, 2016) and pairwise comparisons were further highlighted using ggplot2 (Wickham, 2016). Finally, we used inverse PCA to plot theoretical fruit shapes from PC1 and PC3 using python libraries NumPy (Harris et al., 2020), Matplotlib (Hunter, 2007), scikit-learn (Pedregosa et al., 2011), and seaborn (Waskom, 2021), plotting one value per genetically unique accession, with the accession coloured by aspect ratio.

## RESULTS

In this study, we used a comprehensive pseudo-landmark approach paired with traditional linear measurements to quantify variation in fruit shape and size based on 5724 images of diverse apples. The fruit used in this study were sampled from 743 trees, representing 534 unique accessions from Canada’s Apple Biodiversity Collection.

We used a Spearman’s rank correlation to identify relationships between traditional linear measurements and the first three PCs calculated based on pseudo-landmarking (Figure S1). After adjusting for multiple-testing, 20 of the 45 correlations were significant, with 15 of the 20 significant correlations being positive. The complete set of results are provided in Table S2. Fruit width, length, area, and harvest weight were all significantly positively correlated (*p* < 1 × 10^−15^). The weakest correlation was for fruit width and length (ρ = 0.754; Figure 2a) and strongest for fruit area and harvest weight (ρ = 0.949; Figure 2f). Area had a stronger correlation with width (ρ = 0.940; Figure 2b) than it did with length (ρ = 0.926; Figure 2c). Similarly, weight also had a stronger correlation with width (ρ = 0.908; Figure 2e) than length (ρ = 0.860; Figure 2d). Release year was not significantly correlated with any measurement.

**Figure 2.**
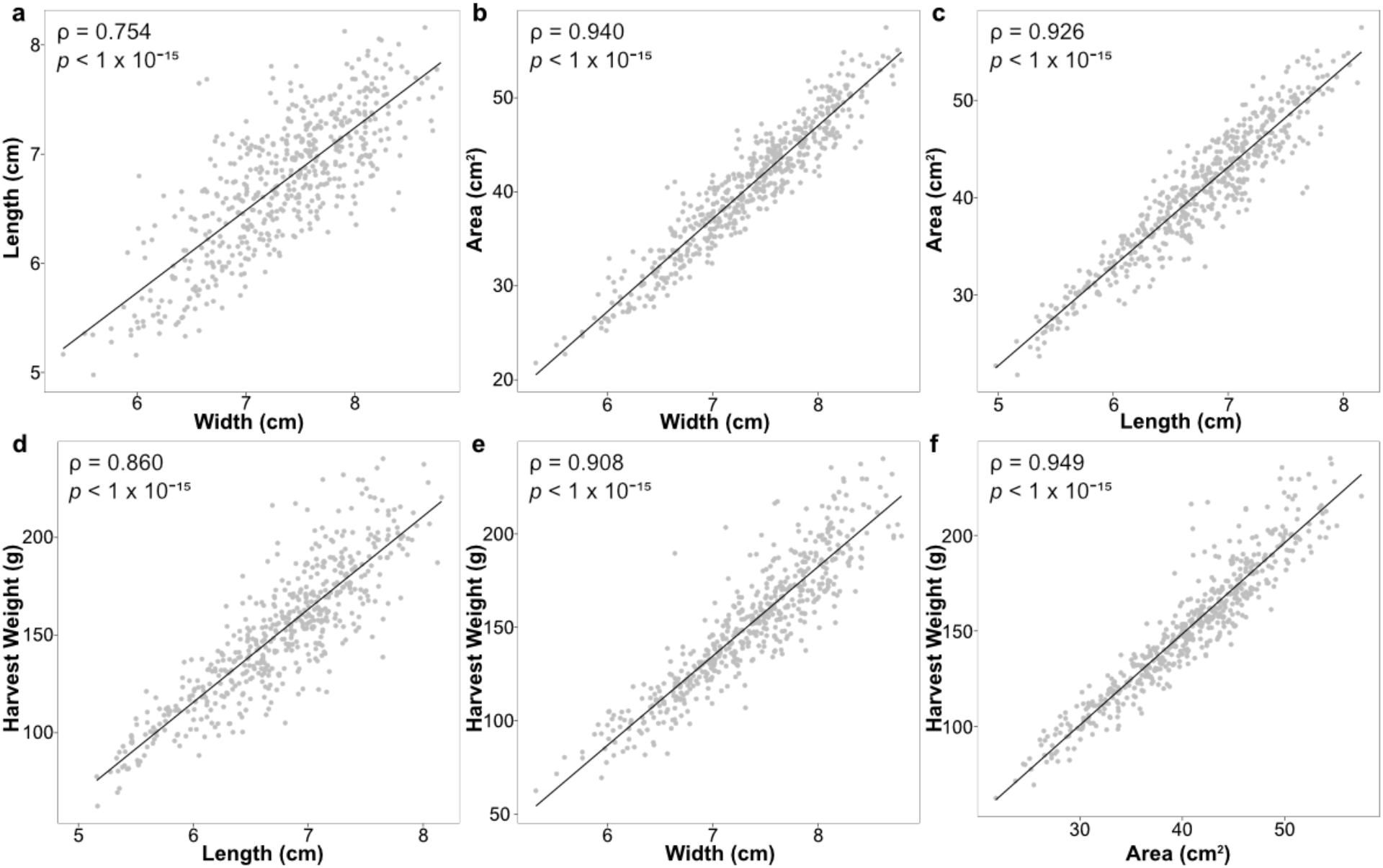
Correlations of measurements from 534 apple accessions for length, width, area, and harvest weight. Spearman’s rank correlation coefficient (ρ) and Bonferroni-correct (based on the total 45 comparisons) associated *p*-value are reported for each correlation.

PCA was performed on pseudo-landmarks to quantify the primary sources of morphological variation in apple. Visual assessment of shapes along the PCs provided insights into the driving forces of the main axes of variation in fruit shape. For example, PC1, the primary axis of variation, explained 22.7% of variation and was primarily related to differences in the aspect ratio (width to length ratio). PC2 (19.0%) was associated with differences in the degree of indentation at the top of the fruit and variation along PC3 (12.4%) represented differences in fruit asymmetry. Combined, the first three PCs explained 54.1% of the variance in fruit shape (Figure S2).

To better visualize variation in fruit shape across all accessions, we generated a morphospace by plotting PC1 against PC3 using one value per unique accession (n = 534). We include PC2 in the visualization of eigenfruit (Figure S2) but not in the morphospace (Figure 3) as the indentation at the top of the fruit is subtle and less likely to be commercially relevant in comparison to the variation captured by PC1 and PC3. For the morphospace, each value is the result of both averaging fruit within a tree and then REML-adjustment based on position within the orchard, reducing the dataset from 5724 individual fruit images to 534 accessions (Figure 3). Each accession was coloured by aspect ratio and plotted on a morphospace which included grey eigen apples, generated using inverse PCA and representing theoretical fruit shapes. It is thus possible to observe both the measured range of morphological variation as well as which theoretical fruit shapes were not measured from the collection.

**Figure 3.**
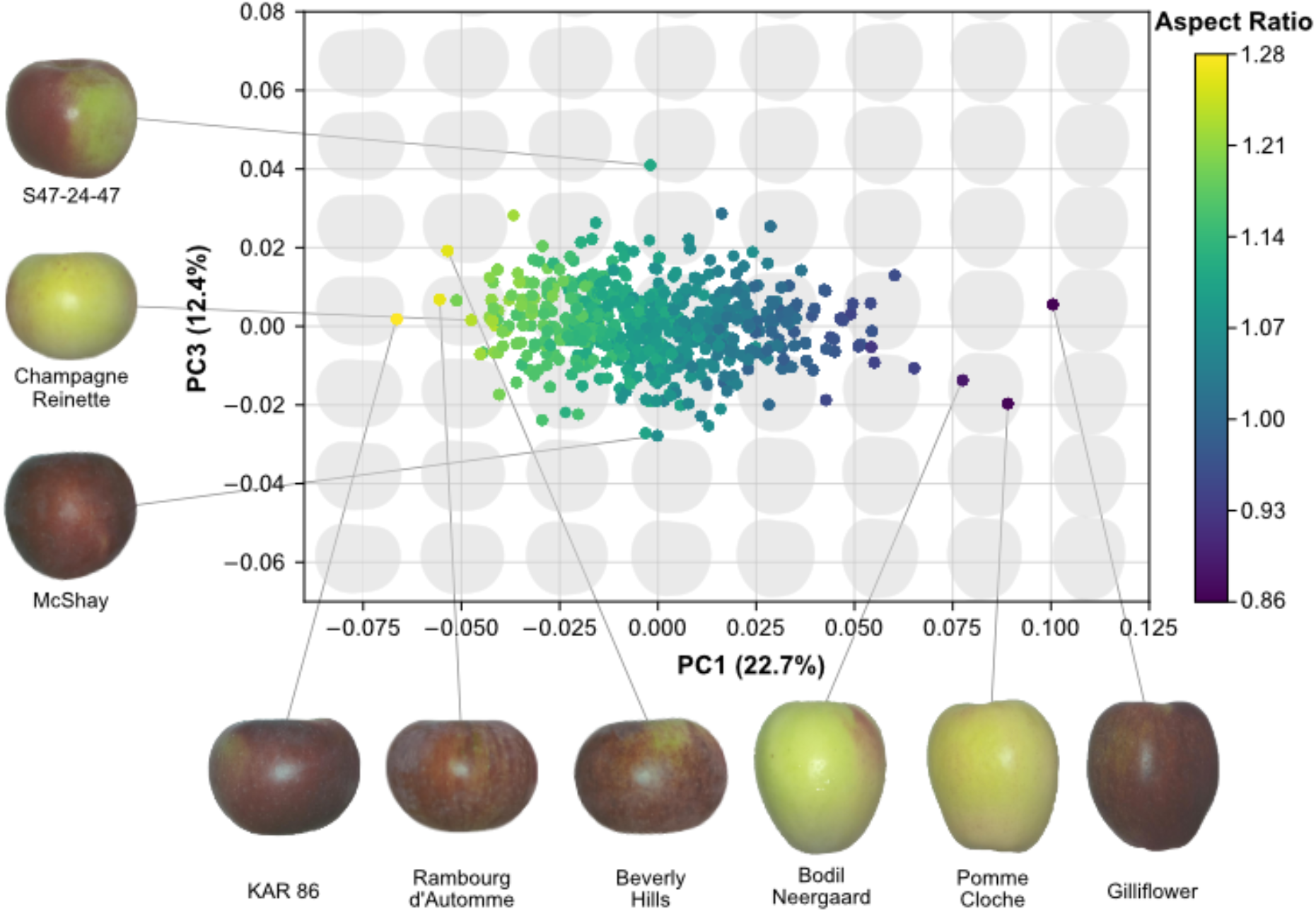
Morphospace of theoretical apples generated using inverse PCA with values for the measured 534 apple accessions indicated along PC1 and PC3. The amount of variation explained by each PC is reported in parentheses and accessions are coloured based on their aspect ratio.

Examining the morphospace reveals a strong relationship between PC1 and aspect ratio, which we quantified using a Spearman’s rank correlation. Out of all 45 pairwise correlations for fruit shape and size evaluated in this study, PC1 and aspect ratio had the strongest correlation (ρ = – 0.964, *p* < 1 × 10^−15^; Figure 4a). Next, we assessed if PC1, the primary source of variation in fruit shape, was also correlated with fruit size, but it was not significantly correlated with fruit area (ρ = 0.126, *p* = 0.162; Figure 4b) or weight (ρ = 0.066, *p* = 1.000; Figure 4c).

**Figure 4.**
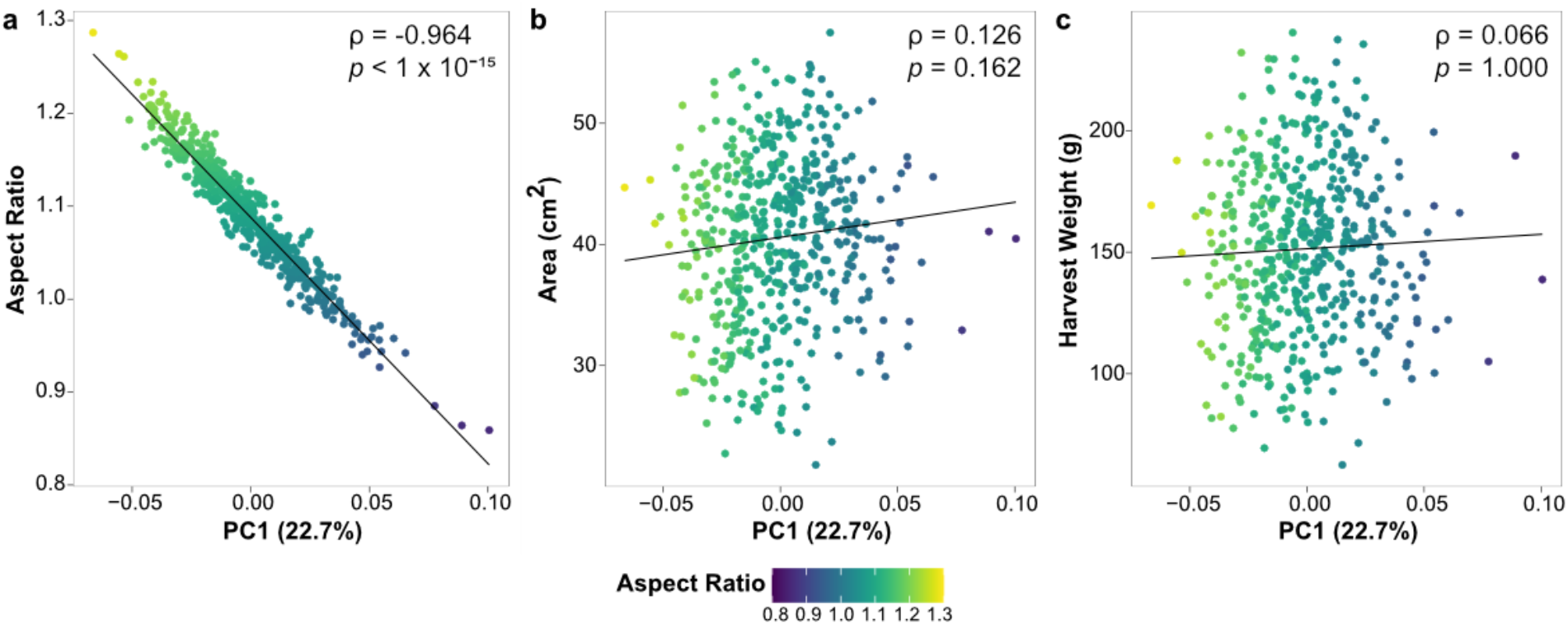
Correlations between PC1, calculated using a pseudo-landmark approach, and (a) aspect ratio, (b) area, and (c) harvest weight across 534 apple accessions. Spearman rank correlation coefficients (ρ) and *p*-values are reported for each correlation.

Taken together, these findings demonstrate that the primary source of variation in fruit shape is the aspect ratio, i.e. the ratio of fruit width to length, and that variation in aspect ratio is independent of fruit size.

## DISCUSSION

The shape and size of apples are critical traits when it comes to consumer preferences and visual assessments of fruit. In this study we quantified morphological variation across Canada’s Apple Biodiversity Collection, one of the world’s largest and most diverse collections of apples (Watts et al., 2021).

We identified aspect ratio as the primary source of variation in fruit shape. Accordingly, accessions with the lowest aspect ratio and highest PC1 values were ‘Gilliflower’, ‘Pomme Cloche’, and ‘Bodil Neergaard’ (Figure 3). These accessions have an elongated, cylindrical shape, often described as oblong. All three are relatively old accessions and are considered heritage apples: ‘Gilliflower’ dates back to 1813 in England, ‘Bodil Neergaard’ to 1850 in Denmark, and ‘Pomme Cloche’ to 1939 in Switzerland (United States Department of Agriculture, Agricultural Research Service, 2024; Watts et al., 2021). While release year was not significantly correlated with any shape or size measurements across our entire data set, the older release dates of these unusually shaped accessions suggest consumers during those eras had a stronger preference for oblong fruit shapes than they do today.

Accessions with the highest aspect ratio and lowest PC1 values were ‘KAR 86’, a breeding line from Canada, ‘Rambourg d’Automme’ from France, and ‘Beverly Hills’ from the United States (Figure 3). These accessions were wide and short, having a flat or oblate shape. Similar to accessions with very high PC1 values, none of these apples are widespread in cultivation, suggesting that that they may offer novel variation in fruit shape for breeding, marketing, or research. For example, accessions with extreme high and low PC1 values could be used for future functional studies of fruit morphology and in validation of candidate genes from previous work. A recent study by Dujak et al. (2024) combined a genome-wide association study with RNA-sequencing to detect differences in the expression of ovate family protein genes on Chromosome 11 by contrasting a flat and an oblong apple variety. The results of our study could enable future work investigating if these candidate genes differ across other apples with extreme variation in fruit shape.

The variation across PC2 primarily captured a change in the indentation at the top of fruit, which reveals the ability of comprehensive morphometric approaches like pseudo-landmarking to capture suble differences in shape that may be otherwise difficult to quantify. Although subtle, future work could explore if there is any evidence of consumer preferences for this morphological trait.

In contrast to the more subtle variation across PC2, PC3 reveal strong differences in fruit asymmetry. The higher PC3 values have a right asymmetrical side like ‘S47-24-47’ from Canada, while the lower PC3 values have a left asymmetrical side like ‘McShay from the United States’ (Watts et al., 2021). Given that the right and left side of the fruit are not biological meaningful and rather an artifact of the positioning of the fruit when photographed, the contrast to both high and low PC3 values are accessions with PC3 values of approximately zero, representing highly symmetrical fruit shapes such as ‘Champagne Reinette’.

While asymmetrical shape may be caused by the internal structure due to seeds and pollination (Brault & Oliveira, 1995; Carisio et al., 2021), extreme asymmetry can deter consumers from choosing those fruits (Kays, 1999). Having more symmetrical fruit aligns with the consumers cosmetic preferences of meeting a certain visual standard (Loebnitz et al., 2015). If a genetic basis to fruit asymmetry could be detected, this would serve as a desirable target for avoidance in breeding in apple cultivars.

Many morphometric studies in plants have studied variation in leaf shape (Chitwood, Rundell, et al., 2016; Chitwood & Otoni, 2017; Migicovsky et al., 2022), although there are also examples of seed shape (Bouby et al., 2024; Terral et al., 2012) and fruit (Feldmann et al., 2020; Gonzalo et al., 2009; Somogyi et al., 2022). However, we could identify limited examples of comprehensive morphometrics of apple fruit shape (Christodoulou et al., 2018; Currie et al., 2000; Dujak et al., 2023; Keller et al., 2024), particularly across diverse germplasm. A previous study examined a biparental cross (‘Jonathan’ x ‘Golden Delicious’) to measure fruit weight, length, width, and aspect ratio. There was no significant correlation between size (weight) and shape (aspect ratio) and the work described independent genetic control of these characteristics (Chang et al., 2014). Consistent with these findings, we find fruit size and PC1, which captures primarily aspect ratio, are independent in a much broader, genetically diverse, collection. In addition to PC1, neither PC2 (indentation) or PC3 (asymmetry) were significantly correlated with variation in fruit size, providing further evidence that morphological variation in fruit shape across diverse apples is not size-dependent (allometric).

Previously, we measured leaf shape in Canada’s Apple Biodiversity Collection using many of the same trees and identified aspect ratio of the blade as the primary source of variation (Migicovsky et al., 2018). In this previous study, we used elliptical Fourier descriptors and persistent homology and found that PC1 explained 80.23% and 62.20% of the variance in leaf shape, respectively. However, PC1 was significantly correlated with the width of the leaf, resulting in allometric variation. In contrast to leaf shape, we detect no association between shape and size in fruit from the same trees, suggesting that leaves and fruit differ in their developmental constraints.

Understanding shape variation in apples has the potential to impact the future of plant breeding by providing valuable insights that can guide more efficient and targeted breeding efforts. Shape is repeatedly cited as a defining characteristic that consumers consider when selecting apples (Bolos et al., 2021; Normann et al., 2019). Fruit size is also critical, with medium to large apples often chosen over smaller fruit, with the preferred size being 7.4 - 7.7 (± 1.0) cm in diameter (Hampson et al., 2002; Hampson & Quamme, 2000; Jesionkowska & Konopacka, 2006). The independence of fruit shape and size observed in our study suggests that it should be possible to select for differences in fruit shape without significantly impacting fruit size, which has important implications for consumer approval and purchasing decisions.

Ultimately, this study provides new insights into the independence of fruit shape and size in apple as well as a foundation for understanding the genetic factors that contribute to biological variation in apple shape. This research not only improves our understanding of fruit morphology but also assists plant breeders in selecting for apples with novel and desirable traits.

## Acknowledgements

This work was supported by funding from the Canada Research Chairs program, a Natural Sciences and Engineering Research Council of Canada (NSERC) Discovery Grant, the Acadia University Research Fund, and US National Science Foundation Plant Genome Research Program awards (IOS-2310355, IOS-2310356, and IOS-2310357).

## Author contributions

ZM conceived of the initial idea for this study with input from DHC and SM. ML aided in methods development and image processing. KD analyzed the data with input from DHC and ZM. SM and ZM acquired funding for this study. KD and ZM wrote the manuscript, which all authors read, edited, and approved, prior to publication.

## Data availability statement

The original images of fruit used for this study as well as the converted binary images are available from the Dryad Digital Repository: URL TO COME. All data and code used in this study can be found on GitHub: https://github.com/kylidevi/apple_shape

Additional supporting information may be found online in the Supporting Information section at the end of the article.

## Online Supporting Information

**Figure S1**. Pairplot of all linear measurements, harvest weight, release year, and calculated PCs. Each measurement is displayed as scatterplots for the comparison of relationships between each pair of measurements. Density plots along the diagonal represent the distribution of each individual traits.

**Figure S2**. Eigen representation of the first three principal components (PC) representing a combined total of 54.0% of variation in fruit shape, with mean fruit as well as ± 2, 4, and 6σ.

**Table S1**. Data table for all coordinates from each accession along with their calculated linear measurements, averaged linear measurements, harvest weight and their release year

**Table S2**. Data table for Spearman’s rank correlation results between all 45 pairwise correlations. Both ρ and *p*-values (after Bonferroni correction) are reported.

