## Supplementary figures and images for "Differences in apple fruit shape are independent of fruit size"

### Figure S1

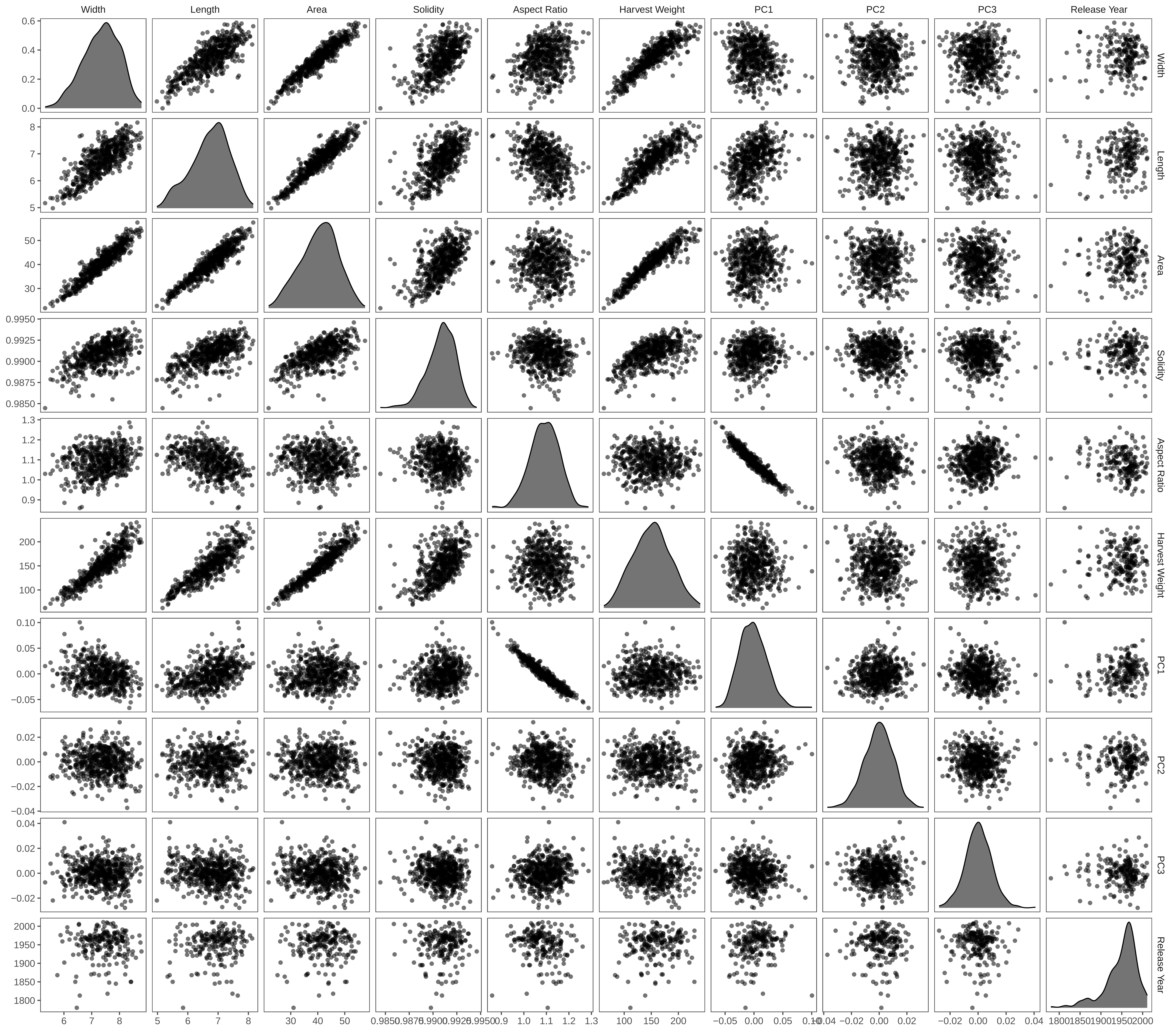

### Figure S2

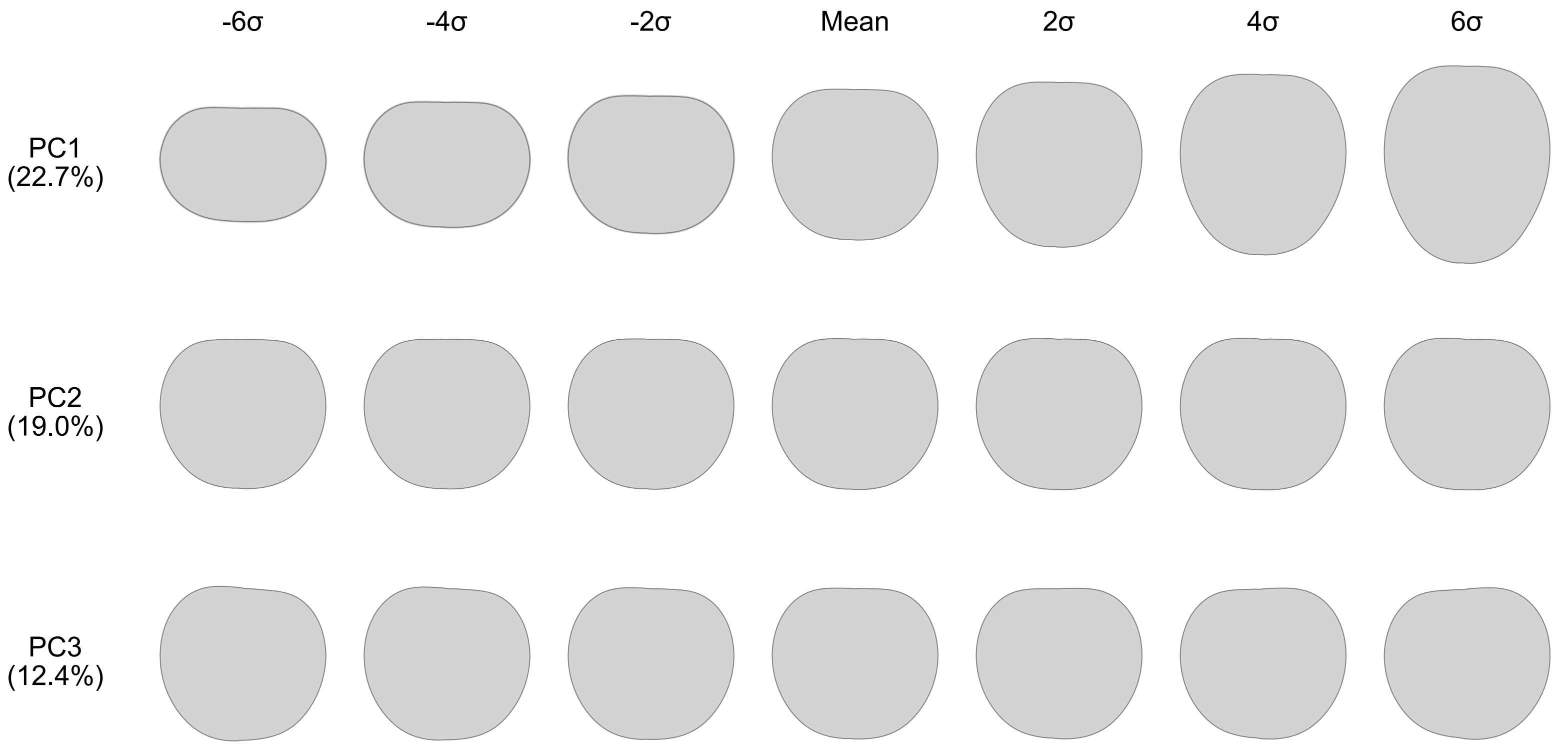
